# Trajectory-centric Framework TrajAtlas reveals multi-scale differentiation heterogeneity among cells, genes, and gene module in osteogenesis

**DOI:** 10.1101/2024.05.28.596174

**Authors:** Litian Han, Yaoting Ji, Yiqian Yu, Yueqi Ni, Hao Zeng, Xiaoxin Zhang, Huan Liu, Yufeng Zhang

**Affiliations:** The State Key Laboratory Breeding Base of Basic Science of Stomatology (Hubei-MOST) and Key Laboratory of Oral Biomedicine, Ministry of Education, School & Hospital of Stomatology, Wuhan University, Wuhan, 430079, Hubei, PR China; Medical Research Institute, School of Medicine, Wuhan University, Wuhan, 430071, Hubei, PR China

## Abstract

Osteoblast differentiation is crucial for bone formation and maintaining skeletal integrity. Although it is now understood that this process exhibits significant heterogeneity across developmental stages and tissue microenvironments, the underlying mechanisms remain largely unexplored. In the present study, we introduce **TrajAtlas**, a comprehensive framework that addresses this gap in knowledge. **TrajAtlas** comprises four modules: a reference atlas (**Differentiation Atlas)**, a differentiation model (**Differentiation Model)**, a tool for differential pseudotime analysis (**TrajDiff)**, and a method for pseudotemporal gene module detection (**TRAVMap)**. By leveraging single-cell technologies, **TrajAtlas** offers a systematic approach to exploring the multi-scale heterogeneity among cells, genes, and gene modules within population-level trajectories across diverse tissues and age groups. We systematically investigate the impact of age and injury on osteogenesis, providing new insights into osteoporosis and bone regeneration. In conclusion, our comprehensive framework offers novel insights into osteogenesis and provides a valuable resource for understanding the complexities of bone formation.

**Author Summary:** Osteoblasts, the cells responsible for bone formation, can originate from various cellular sources. However, it’s unclear how different progenitor cells differentiate into osteoblasts, and how this process is influenced by factors such as age and tissue location. This knowledge gap stems from the lack of comprehensive databases and tools to decipher the differentiation process. In this study, we introduce TrajAtlas, a comprehensive framework designed to bridge this gap. To explore the cellular origins of osteoblasts, we constructed an atlas centered on osteogenesis. To answer how progenitor cells differentiate to osteoblasts, we developed a model that reveals the dynamic regulatory landscape during this process. To elucidate the influence of age and tissue location on differentiation, we built a tool for differential analysis. Furthermore, to identify conserved patterns of differentiation, we developed an approach to detect pseudotemporal gene modules. We validated the effectiveness of this framework by applying it to more datasets, unveiling novel cell states associated with injury. Notably, this framework focuses on dynamic processes, with the potential for broader applications in studying cell differentiation and complementing cell-centric analyses.

## Introduction

Osteoblasts, or bone-forming cells, play a critical role in the dynamic processes of bone formation^1 2^, remodeling^2 3^, and regeneration^4^. Due to their limited lifespan, osteoblasts require constant replenishment by osteoprogenitor cells (OPCs)^5^. This process, known as osteoblast differentiation^5^, occurs in various tissues and different ages in response to various stimuli such as development and regeneration^5^. Across these tissues and age groups, diverse osteoprogenitors contributing to this process have been identified^5 6^. In this respect, recent studies have sequentially identified bone marrow stromal cells expressing markers *Lepr*^5 6 7^, *Grem1*^5^, and *Cdh2*^5^ as osteoprogenitors in long bone. Besides, chondrocytes, expressing *Pthlh*^5 6 8^, *Foxa2*^9^, and *Cd168*^8^, were also identified as cellular sources of osteoblast in long bone. There has been a long-standing debate about how much these osteoprogenitors overlap and how they differently contribute to osteogenesis^5 6^.

The ambiguous definition of osteoprogenitor cells has largely hindered our understanding of osteoblast differentiation. Multiple studies have proposed distinct pathways for osteogenesis in long bones, resulting in a fragmented view of the process^5 6 8 10 11^. While efforts to synthesize these findings into a unified framework have been undertaken^12^, they failed to illuminate the intricacies of osteoblast differentiation. The current classification system, which dichotomizes osteogenesis into endochondral and intramembranous based on the presence of a cartilaginous template^13^, overlooks the true cellular origins of osteoblasts. This oversimplification emphasizes the need for a novel model that incorporates the origin of osteoblasts, thereby providing a more accurate representation of osteoblast differentiation and addressing the existing knowledge gaps.

During cell differentiation, trajectories with distinct gene expression and transcription factor activities emerged, leading to variations in the final cell phenotype^14^. We refer to this phenomenon as differentiation heterogeneity^15^. Several factors contribute to this heterogeneity, including age^16 17^, tissue location^5 6^, and injury^7^. In this context, studies have shown that the expression of certain genes, such as *Maf*, declines with age^16^. Conversely, under injury conditions in long bones, the Wnt signaling pathway is upregulated to promote bone regeneration^7^. Differentiation heterogeneity is intricately linked to age-associated bone disorders, such as osteoporosis^18^, highlighting its significance in the field of bone health. A systematic exploration of this differentiation heterogeneity holds the promise of uncovering new drug targets and propelling forward the fields of drug development^19^ and biomedical engineering^20^, offering innovative solutions to bone-related ailments.

In recent years, single-cell technologies have revealed cellular states with unprecedented detail, providing detailed insights that facilitate the reconstruction of differentiation processes^21^. Most single-cell atlases^22 23^ and tools^24 25^ have focused on cellular state, thereby enabling the identification of cell-type-specific genes, pathways, and gene modules. However, current cell-centric analysis do not fully address the complexities of differentiation heterogeneity. While there are tools designed for trajectory inference and pseudotime analysis, which offer some insights into the temporal aspects of cellular differentiation^26 27 28^, they fall short of providing a comprehensive exploration of cellular dynamics. Specifically, a critical gap exists in the repertoire of available tools. These tools cannot to systematically investigate cells, genes, and gene modules across the diverse spectrum of differentiation trajectories in a dynamic manner. This limitation underscores the need for more sophisticated analytical frameworks that can integrate and interpret the vast and intricate data generated by single-cell technologies, offering a more nuanced and holistic view of cellular differentiation processes.

To address the complexity and heterogeneity inherent in osteoblast differentiation, our study introduces **TrajAtlas**, a trajectory-centric framework specifically designed to navigate and elucidate these challenges. **TrajAtlas** encompasses a reference atlas for osteoblast differentiation coupled with a sophisticated seven-layer hierarchical annotation system, allowing for the detailed modeling of differentiation pathways from diverse osteoprogenitor cells to mature osteoblasts. It features advanced tools for dissecting local variations along trajectories, facilitating the identification of differentially expressed genes during differentiation, and uncovering trajectory-related gene modules to infer their activity within these paths. By leveraging **TrajAtlas** for an in-depth analysis of cellular, genetic, and gene module heterogeneity in large-scale trajectories, particularly focusing on the influences of age and injury on bone formation, our framework unveils novel insights into osteoporosis and bone regeneration, promising significant advancements in bone research and therapeutic strategies.

## Results

### Overview of TrajAtlas and its implementation for osteogenesis

**TrajAtlas** is composed of four modules, enabling the unraveling of differentiation heterogeneity in a multi-scale manner. In the first module, **Differentiation Atlas**, we integrated trajectories spanning a wide range of tissues and ages to establish a reference atlas for osteoblast differentiation, encompassing 110 samples and 272,369 cells. We provided a seven-level hierarchical annotation system to reconcile conflicting osteogenesis-related cell types annotated across different studies. Through mapping experimentally validated osteoprogenitor cells to our atlas, we can delve deeper into exploring the heterogeneity of osteoprogenitor cells (Figure 1a). In the second module, we introduced the **Differentiation Model**, which aims to reconstruct the osteoblast differentiation process. The **Differentiation Model** leverages four trajectories groups to effectively represent differentiation from various osteoprogenitor cells, offering valuable insights into the key signaling pathways and gene regulatory networks underlying this intricate process (Figure 1b). In the third module, **TrajDiff**, we presented a tool designed to detect covariate-related differential cell abundance and genes along the differentiation process across multiple trajectories, enabling assessment of the impact of covariates such as age and tissue locations on gene expression during different stages of osteoblast differentiation (Figure 1c). In the fourth module, **TRAVMap** was introduced, offering innovative methods for identifying conserved variations in trajectories, referred to as Trajectory-related Replicable Axes of Variation (TRAVs). Through TRAVs, trajectory-related gene modules were pinpointed, elucidating their activities within large-scale trajectories (Figure 1d).

**Figure 1.**
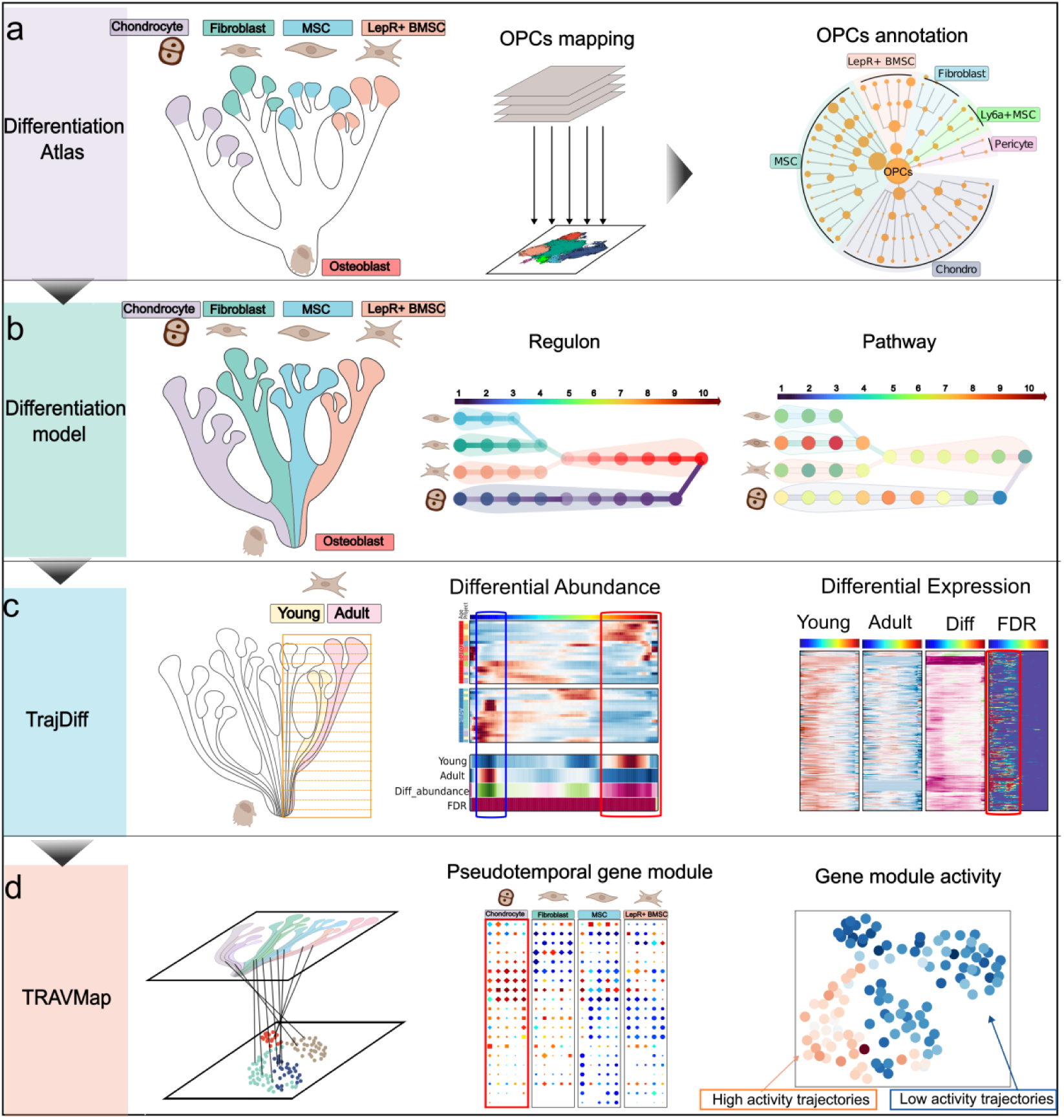
Overview of TrajAtlas: A framework designed to unravel osteogenesis heterogeneity in a multi-scale manner. **a, Differentiation Atlas** integrates trajectories spanning various tissues and continuous age groups to construct a differentiation atlas aimed at identifying various osteogenic precursor cells (OPCs). **b**, **Differentiation Mode**l reconstructs the osteoblast differentiation process, unveiling key genes and transcription factors associated with OsteoProgenitor Cell-Specific Trajectory (OPCST). **c, TrajDiff** detects covariates-related differential cell abundance and gene expression along the differentiation process across multiple trajectories **d**, **TRAVMap** module identifies trajectory-related gene modules and infers gene module activity across large-scale trajectories.

### The osteogenic differentiation atlas reveals the heterogeneity of osteoprogenitor cells

To construct a comprehensive map of osteoblast differentiation, we combined 26 datasets into the **Differentiation Atlas**, encompassing a total of 272,369 cells (Figure 2a). These datasets originated from three primary osteogenic tissues (head, limb bud, and long bone^5 6^) across various age groups, ranging from embryo to old age (see Methods). With a focus on the differentiation process, we filtered out cells irrelevant to osteogenesis (Figure 2a, Supplementary Figure 1a). Next, we employed **scANVI**^29^ to integrate the datasets, preserving biological variations while removing batch effects within the atlas (Supplementary Figure 2a-d, Methods). Subsequently, we implemented a multi-level clustering method to visualize the hierarchical organization of cell populations (Methods). We manually annotated the first three levels of clusters based on previous studies^30^ and then utilized marker genes to distinguish finer cell states (Figure 2c, Supplementary notes 2). Furthermore, we harmonized findings from previous studies within the atlas and provided detailed descriptions of most clusters in the supplementary table and online websites (Supplementary table 2, Data availability).

**Figure 2.**
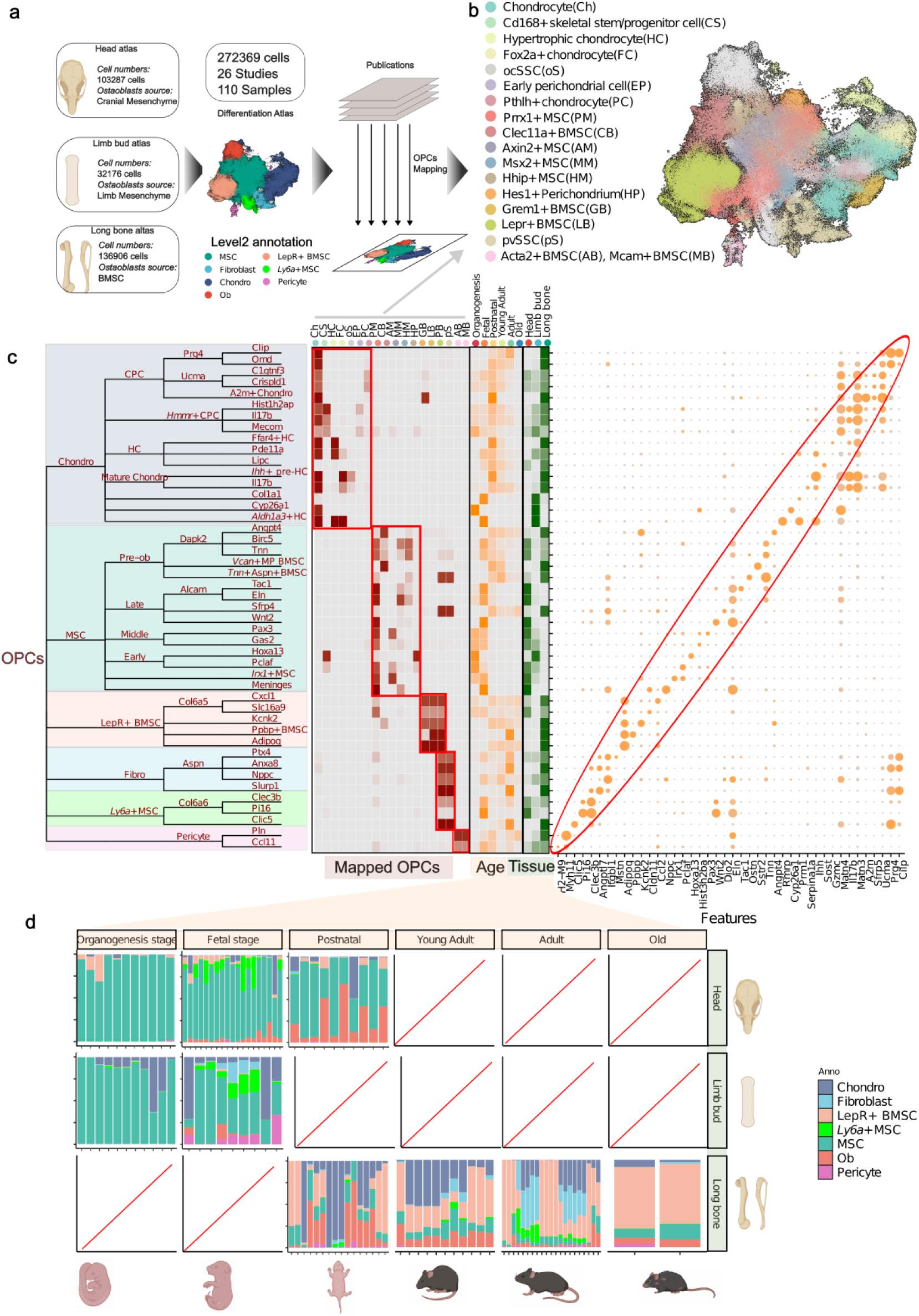
Differentiation atlas reveals the heterogeneity of Osteoprogenitor cells. **a**, A schematic for differentiation atlas. **b**, UMAP visualization of experimentally validated osteoprogenitors mapped to **Differentiation Atlas**. (**c**)A hierarchical tree of clusters of Differentiation **Atlas**. The first five levels with up to 49 clusters are presented, emphasizing the diverse nature of osteoprogenitors across various tissues and age groups. The left heatmap (red) depicts the overlapping of experimentally validated osteoprogenitors with clusters in the lowest tree level in the **Differetiation Atlas.** The middle heatmap (orange) depicts the relative percentage contribution of each cluster at the lowest tree level to the age group. The right heatmap (dark green) illustrates the relative percentage contribution of each cluster to the tissue origin group. In the right panel, dotplot displays marker genes (Methods) at level 5. **d,** Barplots illustrates the proportions of cell types annotated with level-2 annotation across 65 samples with different tissue and age groups.

We found that osteoprogenitor cells exhibited diverse distributions^6^ and expressed distinct characteristic markers (Figure 2c), as reported in previous studies^5 6^. To assess the osteogenic potential of cells within our atlas, we mapped experimentally validated osteoprogenitors based on previously described marker genes and tissue locations (Figure 2a). Notably, the UMAP and the force-directed graph revealed diverse osteoprogenitor populations arranged radially around osteoblasts, suggesting a potential transition between osteoprogenitor cells and osteoblasts (Figure 2a, Figure 3a). Utilizing our level-2 annotation system, we categorized these osteoprogenitor cellls into six major cell types: chondrocytes, mesenchymal stem cells (MSCs), *Ly6a*+ MSCs, LepR+ bone marrow stromal cells (LepR+ BMSCs), fibroblasts, and pericytes (Figure 2c).

**Figure 3.**
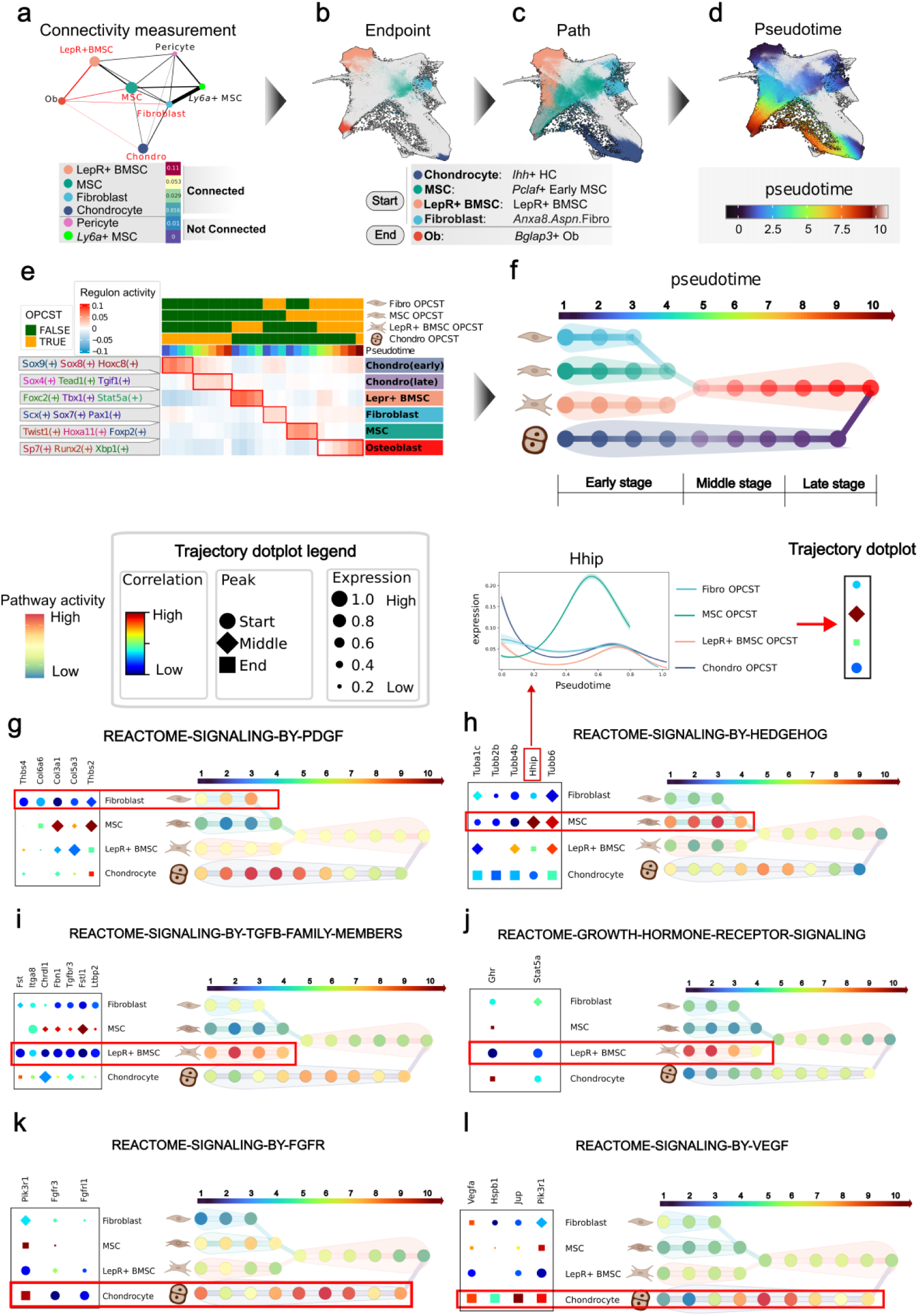
Differentiation Model provides a comprehensive understanding of osteogenic differentiation. **a,** PAGA captures the cell transition of four osteoprogenitor clusters to osteoblasts through coarse-grained connectivity structures. **b,** Force-directed graph of differentiation atlas colored by the endpoint of the four osteoprogenitor clusters. **c,** Force-directed graph of differentiation atlas colored by differentiation path from the four osteoprogenitor cell-specific trajectories (OPCST). **d**, Force-directed graph of differentiation atlas colored by common pseudotime of the four OPCST. **e** Heatmap showing the activity of six regulon clusters (row) across different pseudotime bins of four OPCST (column). **f**, **Differentiation Model** reconstructs the differentiation process of osteoblast. Each node represents a pseudotime bin of OPCSTs, and colors represent regulon clusters in (**e).** pseudotime bins with similar scANVI representation are merged. **g-l**, Activity of key pathway across four OPCSTs. The right panel depicts a trajectory dotplot of genes in the pathway, where the colors represent the correlation coefficient between gene expression and pseudotime; the size of the dots represents gene expression along the pseudotime; the shape indicates the pseudotime where maximum expression occurs. The left panel illustrates the pathway activity across four OPCSTs. The color represents pathway activity.

Chondrocytes, a major cellular source of osteoblasts in the growth plate^31^ were primarily located in the limb bud and long bone datasets within our atlas (Figure 2c,d). Chondrocytes can be further refined into four distinct clusters, all of which are reported to be osteoprogenitor cells with different spatial-temporal distributions. Chondrocyte progenitor cells (CPCs) typically express *Grem1*^30^ and *Pthlh*^32^, which are related to osteogenesis in the resting zone of the growth plate^5 32^(Figure 2c, Supplementary Figure 5b). *Hmmr*+ CPCs^8^ represent proliferating osteoprogenitors in the articular cartilage and the growth plate (Figure 2c, Supplementary Figure 5a). Hypertrophic chondrocytes (HCs)^5 33^ marked by *Col10a1* can directly differentiate into osteoblasts (Figure 2c). Mature chondrocytes expressing *Foxa2*^9^ may contain long-term osteoprogenitors involved in growth plate regeneration (Figure 2c, Supplementary Figure 5c). Interestingly, the limb bud predominantly contained *Hmmr*+ CPCs, while the head exhibited a higher proportion of CPCs (Figure 2c). This distribution highlighted the potential influence of tissue origin and developmental stage on these subpopulations.

MSCs, characterized by the expression of *Prrx1*^34^, comprised the vast majority of the cell population in the head and limb bud datasets (Figure 2c,d). As age increases, the cellular population of MSCs declines, and their cellular states undergo great variation^34^ (Figure 2c,d, Supplementary Figure 6a,b). We annotated these states according to their prominent age as follows: Early mesenchymal stem cells (MSCs), predominant during the organogenesis stage (E8.5-E14), manifested the early perichondrial marker *Hes1*^35^ (Figure 2c); Middle MSCs, the primary cell population in both organogenesis and fetal stage (E14.5-E18.5), specifically exhibited the suture mesenchyme marker *Axin2*^36^ (Figure 2c, Supplementary Figure 5d); Late MSCs, the major cell population in the postnatal stage (P0-P30), highly expressed *Msx2*^37^ and may represent osteoprogenitors in specific craniofacial regions (Figure 2c). Besides, we found an MSC-like cell state that highly expressed adult stem cell maker *Ly6a* (stem cell antigen-6) (Supplementary figure 5e). This population was notably present during the fetal stage in the head and limb regions (Figure 2d), as described in previous studies^34 38^. Consequently, we termed this cluster *Ly6a*+ MSCs^38^.

LepR+ BMSC cells, well-known for their roles in maintaining the bone marrow vasculature and regulating hematopoiesis^5 6 7^, are multipotent cells residing within long bones^39^. Our atlas revealed LepR+ BMSCs in long bones only after the postnatal stage, with their population expanding as the organism ages (Figure 2d). Interestingly, *Cxcl12*+ BMSC, *Grem1*+ BMSC, and *Cdh2*+ BMSC identified in previous studies^5 6 7^ all demonstrated significant overlap with LepR+ BMSC in our atlas (Figure 2b).

Fibroblasts, characterized by high expression of *S100a4*^30^, were primarily observed in the adult stage (3M-12M) of long bones (Figure 2d). We found that they expressed periosteum maker, such as *Postn*^40^, indicating their potential as a source of osteoblasts (Figure 2c, Supplementary Figure 11l,m). Pericytes, characterized by high expression of *Acta2*^30^ (Figure 2c), constituted a minor population within our atlas (Figure 2d).

Our observations suggest that age is likely the most significant factor influencing the cellular states of osteoprogenitors (Supplementary Figure 6a,b). To further investigate this, we performed a gene-level analysis (Methods). Interestingly, we found that most genes displayed consistent age-related effects across all osteoprogenitor types (Supplementary Figure 6c,d). For instance, genes associated with bone formation, such as *Bmp2*, *Chrdl1*, and *Itgb3*, were upregulated with increasing age across all osteoprogenitors, while cell-cycle-related genes like *Cdk8* and *Parp1* were downregulated (Supplementary Figure 6c). GSEA enrichment results indicated that pathways related to oxidative stress, hypoxia response, and lipid synthesis were upregulated across all osteoprogenitors with increasing age. Conversely, cell cycle activity, mRNA splicing, the Wnt pathway, and metabolism of glycine, serine, and threonine were downregulated (Supplementary Figure 6e). These findings align with previous study^41^, indicating a conserved effect of age on osteoprogenitors, which potentially linked to bone disorders like osteoporosis.

### Differentiation Model provides a comprehensive understanding of osteogenic differentiation

To understand how various osteoprogenitor cells differentiate into osteoblasts differentially, we established a **Differentiation Model** within the Differentiation Atlas. We observed that different samples shared similar transition processes from specific osteoprogenitors, supporting the grouping of these trajectories together for further analysis (Supplementary Figure 7a-c). To distinguish these grouped trajectories from individual sample trajectories, we termed them OsteoProgenitor Cell-Specific Trajectories (OPCSTs). We first investigated which osteoprogenitor clusters could directly transition into osteoblasts. To achieve this, we employed coarse-grained connectivity structures to capture cell transitions^28^ between osteoprogenitors and osteoblasts (Figure 3a, Supplementary Figure 7a). This approach identified four distinct grouped trajectories: Chondrocyte OPCST, LepR+ BMSC OPCST, Fibroblast OPCST, and MSC OPCST (Figure 3a, Supplementary Figure 7d).

Next, to identify the endpoints of the developmental trajectories, we explore the earliest states of osteoprogenitors (Figure 3b, Supplementary figure 8a-c, Supplementary note 3). To determine a universal indicator of differentiation from various osteoprogenitors, we benchmarked multiple machine learning strategies and ultimately employed an LGBMR Regressor to build common pseudotime (Figure 3d, Supplementary figure 9a-h, Supplementary note 4). To build the final model, we divided the differentiation path based on osteoprogenitors and pseudotime into bins, then merged bins representing similar cell states (Figure 3f, Methods, Supplementary figure 10a-g). Furthermore, we conducted gene regulatory network inference to predict potential transcription factors and infer the activity of key pathways involved in osteogenesis (Figure 3e-l, Supplementary Figure 12a-h, Methods). Additionally, we created a trajectory dotplot to visualize the pseudotemporal expression patterns across multiple trajectories (Figure 3g-l, Supplementary Figure 14a-i, Supplementary Note 5). The inferred trajectory path of the **Differentiation Model** shows significant overlap with the reported lineage tracing cells^7 33^, and most of the inferred transcription factors (82/136) have been demonstrated to be associated with bone formation (Supplementary Figure 12m, Supplementary table 5). Besides, a large portion of differentiated genes (938/1852) have been annotated in bone databases^42 43 44^, such as Phylobone (Supplementary Figure 13m, Methods, Supplementary table 6), further validating the model’s ability to capture biologically relevant information about bone development.

Previous research has described the cellular transition between chondrocytes and osteoblasts^33^. This transition can be well illustrated in our model, represented as Chondrocyte OPCST (Figure 3a). In our model, the Chondrocyte OPCST primarily progressed through three stages: prehypertrophic chondrocytes, hypertrophic chondrocytes, and osteocytes^30 33^ (Figure 3e). In this differentiation process, transcription factors and signaling pathways undergo significant changes. In terms of transcription factors, early differentiation is regulated by *Sox9*(+) and *Sox8*(+), which maintain the chondrocyte fate, while late differentiation is regulated by *Tgif1*(+) and *Sox4*(+), both of which are associated with parathyroid hormone^45, 46^ (Figure 3d, Supplementary Figure 12g,h). As for signaling pathways, PDGF signaling (*Col9a1*, *Col6a3*, *Thbs3*) is highly activated in the early differentiation process, while FGFR signaling (*Fgfr3*, *Fgfrl1*, *Pik3r1*) and VEGF signaling (*Vegfa*, *Jup*, *Hspb1*) are involved in the late differentiation process (Figure 3k,l). These results show a significant reprogramming in chondrocytes during the transition toward osteoblast differentiation.

In long bones, LepR+ BMSC cells are typically dormant but become osteogenesis-activated in response to injury^7^. Our model divides this process into three stages: LepR+ BMSC cells, pre-osteoblast cells, and osteoblasts (Figure 3a, Supplementary Figure 7d). Like Chondrocyte OPCST, we observed that transcription factors and pathways exhibited cascading changes.

*Foxc2*(+)^47^ and *Stat5a*(+)^48^ regulate the early stage of differentiation (Figure 3d, Supplementary Figure 12e,f), while *Sp7* and *Runx2* regulate the late stage (Figure 3d, Supplementary Figure 12k,m). TGF-β signaling and growth hormone receptor signaling pathway were highly activated at early differenitation, while extranuclear estrogen signaling at late (Figure 3i,j, Supplementary Figure 13l). Apart from LepR+ BMSC, fibroblasts have attracted more and more attention in long-bone regeneration^40^. Our model delineated the Fibroblast OPCST, which illustrates this differentiation process in fibroblasts (Figure 3a-d, Supplementary Figure 7d). Different from LepR+ BMSC, *Pax1*(+)^49^ and *Scx*(+)^50^ regulate the early differentiation, and the PDGF signaling pathway (*Thbs4*, *Thbs2*, *Col6a6*) was highly active during this stage (Figure 3d,g, Supplementary Figure 12a,b).

MSCs are the main cellular source of both limb buds and heads in the embryo. Our model captured this process and depicted it as the MSC OPCST. At early differentiation, *Twist1*(+)^34^ and *Hoxa11*(+)^51^ were transcription factors that regulate this process, and the Hedgehog signaling pathway (*Hhip*, *Tubb6*, *Tubb4b*) exhibited high activity at this stage (Figure 3d,h, Supplementary Figure 12c,d). Interestingly, we found that MSCs differentiated to pre-osteoblasts sharing similar cellular states with pre-osteoblasts in LepR+ OPCST (Supplementary Figure 10d). However, pre-osteoblasts in MSC OPCSTs demonstrated high activity in the Wnt signaling pathway (*Wnt6*, *Wnt10a*, *Wnt2*), whereas pre-osteoblasts in LepR+ BMSC OPCSTs exhibited high activity in the Complement Cascade (*C3*, *C4b*, *Cfh*) (Supplementary Figure 13n-p). This difference suggested that different osteoprogenitors could retain part of their own identity for responding to the microenvironment even after transitioning into similar cell states.

Overall, our **Differentiation Model** reconstructs the osteoblast differentiation process from different osteoprogenitors and provides a systematic view of the diversity in gene regulatory networks and pathway activities during osteoprogenitors differentiation.

### TrajDiff detects covariate-related gene differences across multiple trajectories

It is now understood that throughout the process of differentiation, variations in both cell abundance and gene expression^26 52^ are highly influenced by factors such as age^16^ and tissue origin^14^. In multi-stage differentiation processes like osteoblast differentiation^3^ (Figure 3f), it is pivotal to pinpoint the stage at which differential genes exert their influence. However, current methods^26 52^ fail to infer the differential abundance and expression within specific differentiation stages, thus calling for the development of new tools. In the present study, we introduced **TrajDiff**, a novel tool specifically designed for performing differential pseudotime analysis across multiple samples, aimed at uncovering changes in cell abundance and expression throughout differentiation stages (Figure 4a). Our benchmarking results demonstrated that **TrajDiff** could outperform existing algorithms, such as **Lamian**^26^ and **Condiments**^52^, in various aspects, including the detection of differential abundance (Supplementary Figure 15a-i, Supplementary Note 6) and trend differences of gene expression (Supplementary Figure 16a-e, Supplementary Note 6).

**Figure 4.**
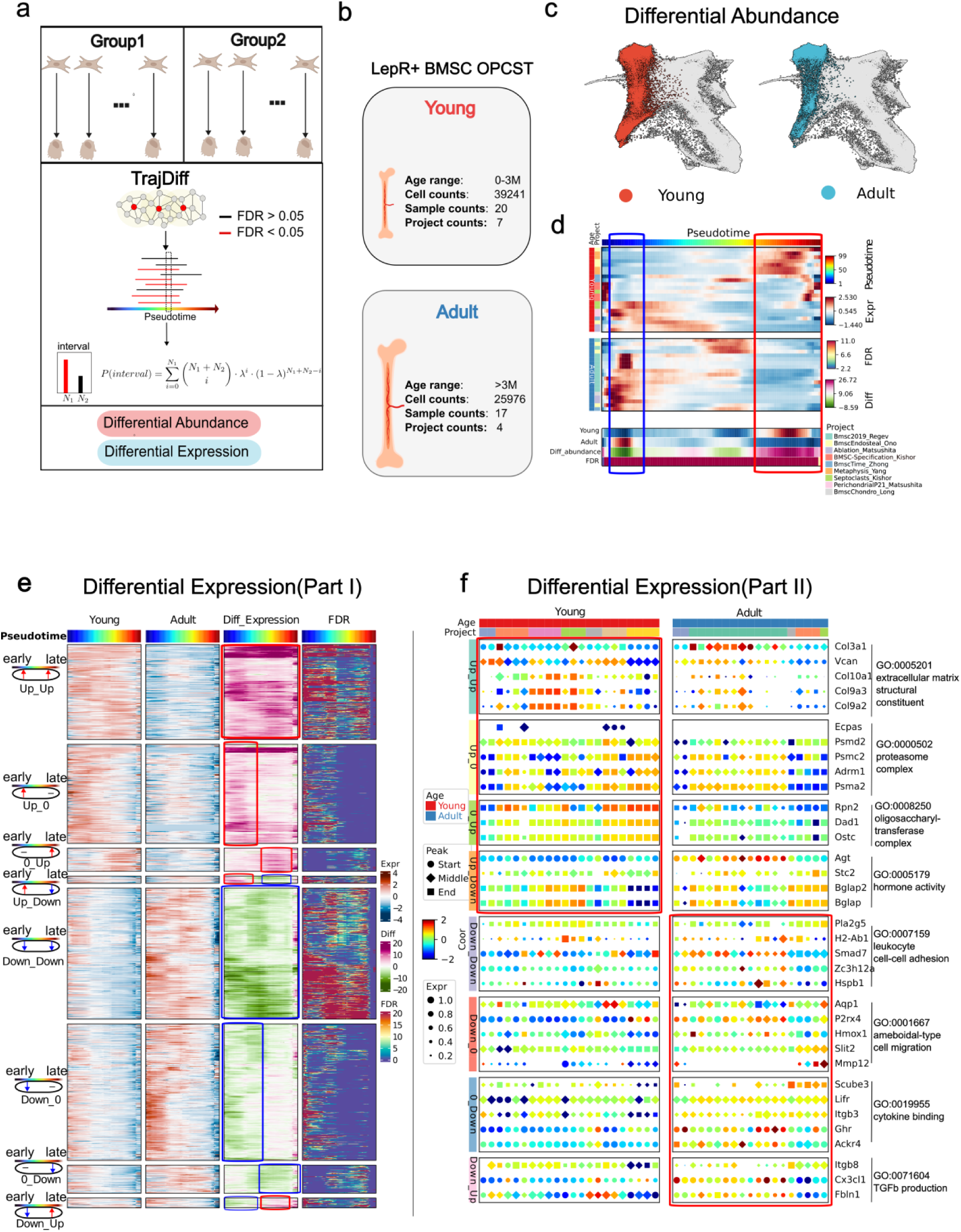
TrajDiff detects covariate-related gene differences across multiple trajectories. **a**, A overview for **TrajDiff**. **b,** Study design for **TrajDiff** analysis. **c**, Force-directed graph visualization illustrates the difference in cell abundance between the Young and Adult groups. **d,** Heatmap illustrates the difference in cell abundance along pseudotime (column) in 37 samples (row) between two groups. The four rows of bottom annotation represent: the mean cell abundance of the two groups (row 1, row 2), differential abundance (row 3), and false discovery rate (FDR) (row 4). **e**, Heatmaps are presented in four vertical panels to illustrate gene expression of the Young group (first panel) and Adult group (second panel), expression differences between two groups (third panel), and FDR (fourth panel). In each panel, rows represent genes, while columns represent pseudotime. Genes are categorized into 8 clusters based on their expression patterns. **f**, Trajectory dotplot illustrates expression of GO-enriched genes from eight gene clusters in (**d**) across 37 trajectories.

Our previous analysis on osteoprogenitors indicated that the states of LepR+ BMSCs and MSCs could change with age (Supplementary Figure 6a-e). Additionally, prior studies have shown significant remodeling of gene expression^53^ and biological function^39 53^ in LepR+ BMSCs during adolescence. This evidence suggested that the differentiation process of LepR+ BMSC may vary across different ages. To explore this age-related heterogeneity, we applied **TrajDiff** to LepR+ BMSC OPCST. Samples younger than 3 months were defined as the “Young” group, while others were assigned to the “Adult” group, based on previous studies^39 53^ (Figure 4b). Differential abundance analysis revealed higher cell density at the late stage (preosteoblast and osteoblast) in the Adult group compared to the Young group, where cell density was higher at the early stage (LepR+ BMSC) (Figure 4c,d). This observation was unlikely to be due to technical sampling bias, as it has been consistently observed across multiple projects (Figure 4d). These results suggested that LepR+ BMSC cell abundance was low during postnatal and adolescent stages, which might explain why LepR+ BMSCs are not the main source of osteoblasts in the long bones of young mice during adolescence^39^.

Using **TrajDiff**, we next performed differential expression analysis to identify genes whose expression varied across differentiation stages. Our study identified 74.2% of the differentially expressed genes reported in the previous study^53^ (Supplementary Figure 18j). While the previous study provided a binary classification of upregulated or downregulated genes^53^, our results offer a more comprehensive understanding by elucidating the precise developmental stages at which these genes exhibit differential expression patterns. We categorized these differentially expressed genes into eight groups based on their expression patterns (Supplementary Figure 18b-j). We noticed that some genes are differentially expressed during the whole differentiation process. For example, genes related to leukocyte cell−cell adhesion, such as *Smad7*, *Hspb1,* and *Pla2g5*, showed higher expression in the Adult group than the Young group during differentiation (“Down_Down”) (Figure 4d,e, Supplementary Figure 18a,g). While some genes are only differentially expressed at the specific differentiation process. Genes involved in ameboidal−type cell migration (*P2rx4*, *Mmp12*, *Slit2*), for instance, were downregulated only at the early stage (“*Down_0*”) (Figure 4d,e, Supplementary Figure 18a,f). Interestingly, we noticed that certain genes were highly expressed in one group during early differentiation, but then became highly expressed in the other group during late differentiation. Genes related to TGF-β production (*Itgb8*, *Cx3cl1*, *Fbln1*) and hormone activity (*Agt*, *Bglap*, *Bglap2*) are both of this type, which suggests the roles of signaling pathways vary depending on age (Figure 4d,e, Supplementary Figure 18a,e,h,i). These findings corroborate previous research that reported enrichment of pathways associated with hematopoiesis, inflammatory responses^53^, and antigen processing in older mice, whereas our results provide higher resolution insights across developmental stages.

We next investigated genes with transient differential expression. These genes, representing a smaller proportion (357 genes) compared to persistently differentially expressed genes (876 genes), exhibited dynamic changes throughout differentiation (Supplementary Figure 17a,c). In the Young group, transiently upregulated genes at the early stage were associated with neuronal activity (*Clstn2*, *Flrt3*, *Shisa9*) (Supplementary Figure 17a,b). Conversely, genes transiently downregulated at the early stage were associated with morphogenesis (*Sox9*, *Lhx1*, *Spg11*) (Supplementary Figure 17a,b). Additionally, *Hes1*, involved in cell differentiation, was downregulated at the middle stage, while *Stk17b*, associated with fibroblast apoptosis, was downregulated at the late stage (Supplementary Figure 17a,b).

In summary, **TrajDiff** offers a distinct advantage over **Lamian** and **Condiments** by identifying differential genes at precise developmental stages. Consequently, it furnishes a comprehensive landscape of age-related gene expression dynamics during the differentiation of LepR+ BMSCs.

### TRAVMap reveals pseudotemporal gene module heterogeneity across all trajectories

During differentiation, gene modules execute functions in a specific temporal order^54^. However, existing gene module identification methods^54^ fail to consider this temporal information. To address this limitation, we developed **TRAVMap**, a method to identify conserved pseudotemporal gene modules across multiple differentiation trajectories (Figure 5a). **TRAVMap** utilizes matrix factorization to identify recurring axes of variation^55^ within these trajectories, termed Trajectory-related Replicable Axes of Variation (TRAVs) (Figure 5a). We found that genes that drive TRAVs show coordinated expression patterns, representing gene modules that function at specific stages of the differentiation process (Supplementary Figure 21d,f,h). As far as we know, this is the first algorithm that identified pseudotemproal gene modules. Furthermore, we found that the identified gene modules are conserved across multiple samples, validating the effectiveness and robustness of our algorithm (Supplementary Figure 21h). In this way, **TRAVMap** identified 226 TRAVs across 121 trajectories (Figure 5b). Furthermore, by leveraging TRAVs and gene expression data, TRAVMap learned representations of trajectories to capture heterogeneity among samples, similar to a tool called scPoli^56^. However, in contrast to scPoli, our approach learns from gene and gene module activities, rendering it more interpretable and enabling a detailed view of gene expression and gene module activity dynamics across population-level trajectories (Supplementary Figure 19a-e, Supplementary Notes 7). Our methods facilitate multi-scale exploration of genes and gene modules in differentiation processes.

**Figure 5.**
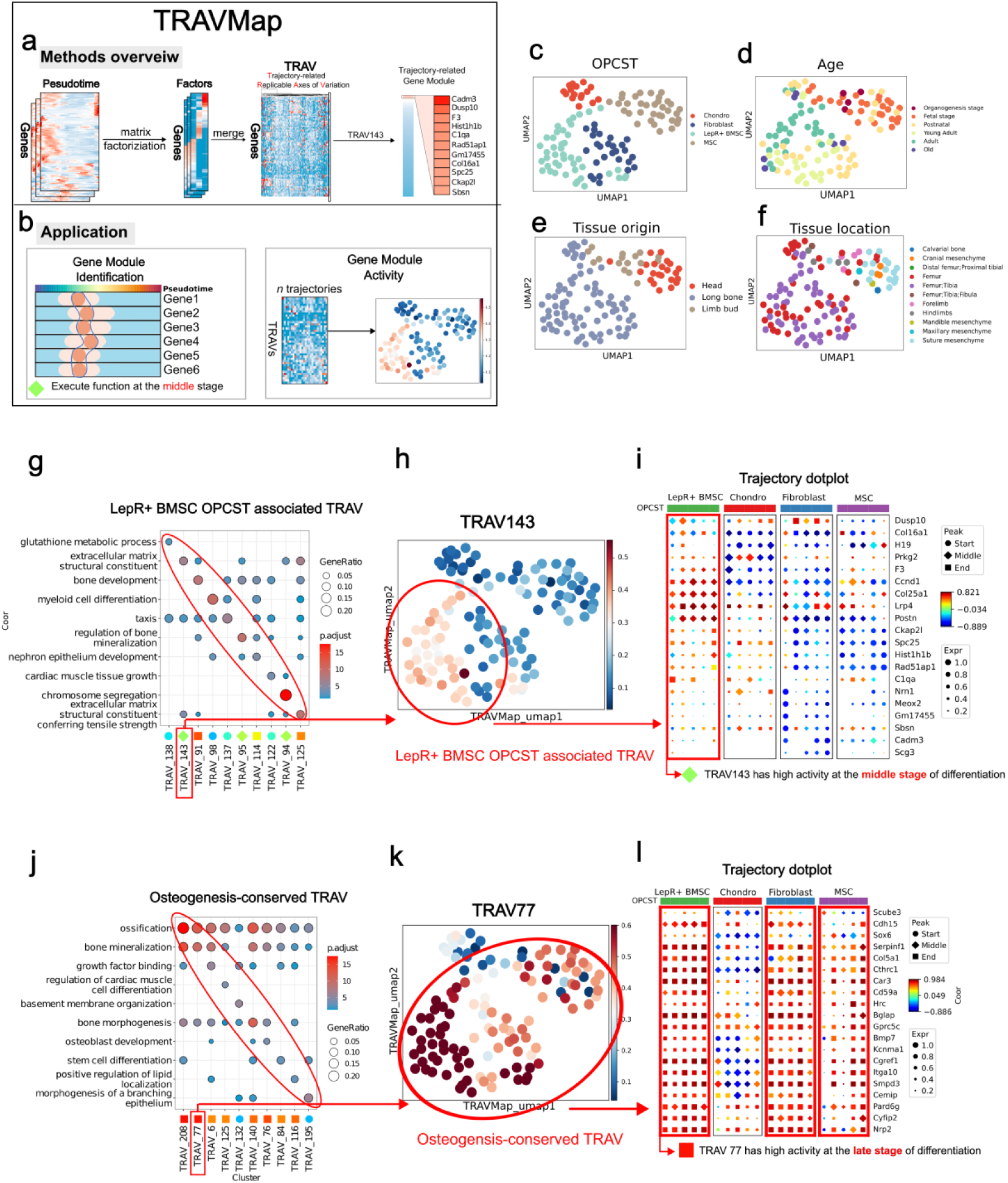
TRAVMap reveals gene module heterogeneity across all trajectories. **a,b**, An overview for **TRAVMap**. **b-e**, Trajectory embeddings visualized with UMAP, colored by OPCST, Age, Tissue origin, and Tissue location. Each point represents a trajectory. **g,j,** GO enrichment of Trajectory related Replicable Axis (TRAV) gene module, visualized with dotplot. The bottom dot annotation represents the differentiation stage at which the gene module executes its function. **h,k,** TRAV activity with UMAP visualization in **(b-e) i,l,** Trajectory dotplot illustrates the gene expression patterns across 20 randomly sampled trajectories from four OPCST.

**TRAVMap** reveals that osteoprogenitor cell identity, age, and tissue origin can all drive heterogeneity in osteoblast differentiation processes (Figure 5c-f, Supplementary Figure 19a-e). Among these factors, the identity of the osteoprogenitor cells contributes the most to this observed heterogeneity. Therefore, we identified dozens of OPCST-associated gene modules. We focused on gene modules in LepR+ BMSC OPCST and found that these gene modules function at specific stages of the differentiation process, representing distinct biological processes. For example, TRAV122 functions at the early differentiation process, enriched genes associated with taxis; while TRAV143 functions at the middle stage of differentiation, enriched genes involved in the structure of the extracellular matrix (Figure 5g-h). To examine the expression pattern of the TRAV143 gene module, we randomly selected five trajectories for each OPCST and visualized them using a trajectory dotplot (Figure 5i). We observed that most of the genes within this module exhibited high expression that peaked at the middle stage of differentiation, specifically in the LepR+ BMSC

OPCST trajectories, whereas they exhibited distinct pseudotemporal expression patterns in trajectories derived from other OPCST (Figure 5i, Supplementary Figure 21h). This confirms that the TRAV143 gene module functioned during the middle stage of differentiation, specifically in the LepR+ BMSC OPCST (Figure 5h,i, Supplementary Figure 21d).

In this way, we identified osteogenesis-conserved gene modules and age-related gene modules. The osteogenesis-conserved gene modules were active across most of the osteoblast differentiation trajectories, representing the conserved genetic programs driving the osteogenic process (Figure 5j, Supplementary Figure 21a). We found that most of these gene modules functioned at the late stage of differentiation, including TRAV208, TRAV77, and TRAV6, which were associated with biological processes like ossification, bone mineralization, and growth factor binding (Figure 5j). The age-related gene modules differentially activate between age groups. One example was TRAV140, which exhibited differential activity across LepR+ BMSC, Fibroblast, and MSC OPCSTs (Supplementary Figure 20a,b). In postnatal and young adult LepR+ OPCST trajectories, TRAV140 showed high activity, and its associated gene module, containing genes like *Smpd3* and *Ano6,* reached peak expression at the late stage (Supplementary Figure 20d,f). However, in adult and old groups, TRAV140 activity was lower, and the gene module exhibited inconsistent expression patterns (Supplementary Figure 20d,g). Notably, a majority of genes (61 out of 100) within the TRAV140 module overlapped with genes identified by **TrajDiff**, further supporting its age-related nature (Supplementary Figure 20e). Gene Ontology analysis suggested that this module was associated with bone morphogenesis and stem cell development, implying potential differences in osteogenesis activity across age groups (Supplementary Figure 20h). Interestingly, the predicted transcription factors regulating TRAV140, *Fosl1*(+)^57^, and *Lef1*(+)^58^, represented promising therapeutic targets for age-related bone disorders like osteoporosis (Supplementary Figure 20i).

In conclusion, **TRAVMap** effectively identified both osteoprogenitor-related and age-related gene modules. This allowed for the creation of a landscape of pseudotemporal gene modules that are active at distinct developmental stages across population-level differentiation trajectories.

### Applying TrajAtlas to an extended atlas elucidates alterations in trajectories under injury conditions

The **Differentiation Atlas** is primarily designed to understand the osteogenesis process in healthy mice, aiming to establish a reference model for normal osteoblast differentiation. However, osteoblast differentiation can be altered or disrupted by various conditions, including injury^7 59^,and disease^59^. To investigate osteogenesis within diverse microenvironments, we projected data from 65 datasets encompassing 319 samples and 781397 cells onto the **Differentiation Atlas** (Figure 6a). This extended atlas incorporated a broader range of tissues (e.g., digit bone, rib, periodontium), encompassed more complex cellular states (injury, heterotopic ossification, disease), and included data from multiple species (human and rat) (Supplementary Figure 22 a-f).

**Figure 6.**
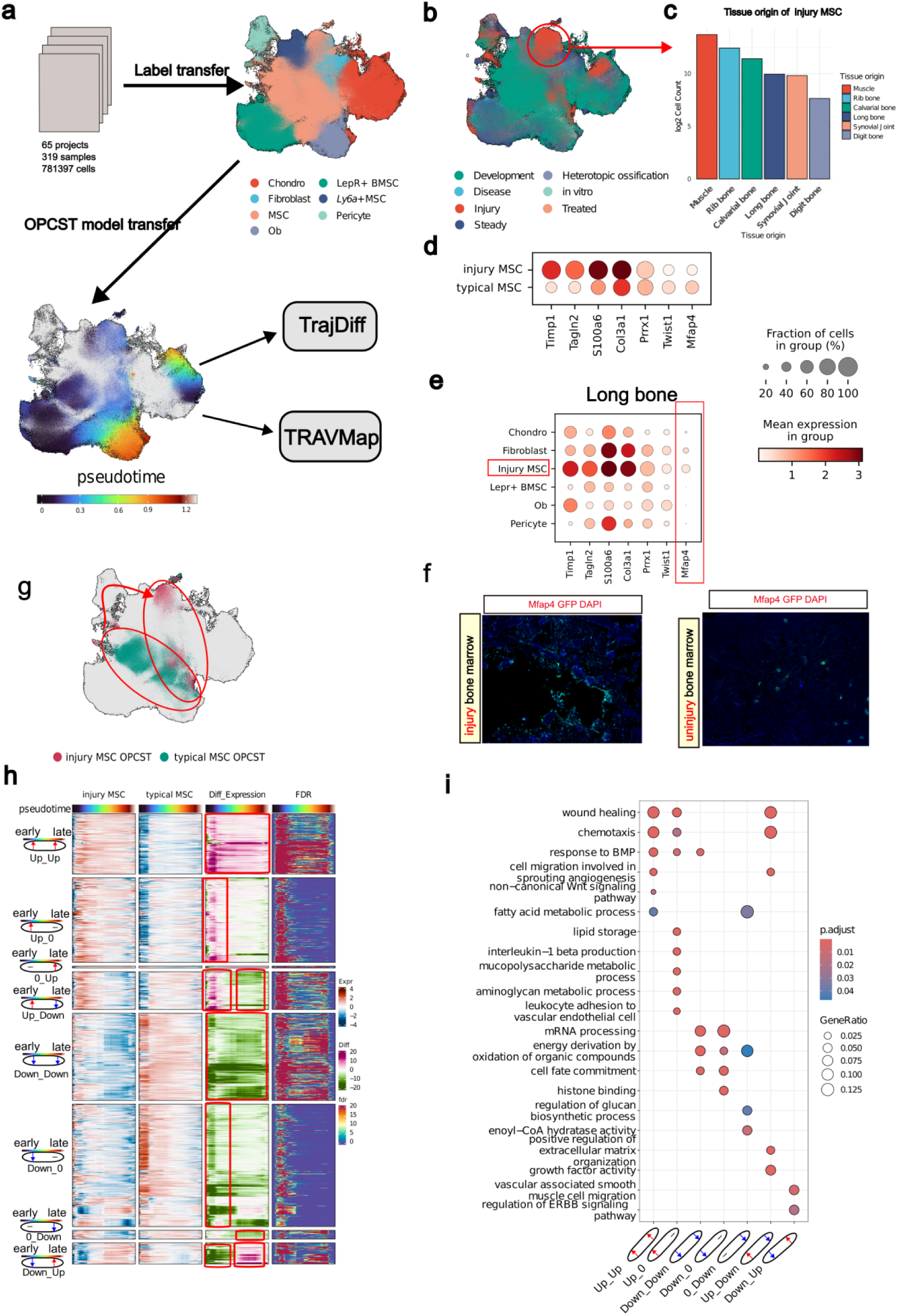
Applying TrajAtlas to an extended atlas elucidates alterations in trajectories under injury conditions. **a**, An overview of extended atlas construction. **b,** UMAP visualization of the extended atlas, colored by stage, highlighting cells (Injury MSC) that group together in the injury state. **c**, Barplot shows the cell count (log_2_ scale) of tissue origin for Injury MSC in (**b**). **d**, Dotplot displays the expression levels of MSC marker genes and differentially expressed genes between typical MSCs and injury MSCs. **e**, Dotplot shows expression levels of marker genes of injury MSC across all cell types present in long bone datasets. f, MFAP4 (green) in 8-week-old femur from injured and uninjured conditions. **g**, UMAP visualization of MSC OPCST between injury and typical states. **h**, Heatmaps are presented in four vertical panels to illustrate gene expression of the injury MSC group (first panel) and typical MSC group (second panel), expression differences between the two groups (third panel), and FDR (fourth panel). **i,** GO enrichment of seven clusters in (**f**), visualized with dotplot.

Following data integration and label transfer, most cells were confidently annotated with level-2 labels (Supplementary Figure 23a-c). However, we observed high uncertainty associated with a specific mesenchymal stem cell state, primarily composed of cells from injury datasets (Figure 6b, Supplementary Figure 23d). Notably, this state, termed “injury MSC”, appeared to originate from various injured tissues such as skeletal muscle, rib bone, and calvarial bone (Figure 6c, Supplementary Figure 23e). This suggests that cells from different tissues may reprogram to an MSC-like state upon injury, reminiscent of blastema formation in salamanders^60^. In salamanders, cells can dedifferentiate and acquire pluripotency to regenerate tissues. Interestingly, in this study, injured MSCs expressed typical MSC marker genes, such as *Prrx1*^34^*, and Twist1*^34^ (Supplementary Figure 23f,g), but also exhibited high expression of genes related to angiogenesis, such as *Tagln2*^61^, which distinguished them from typical MSCs (Figure 6d, Supplementary Figure 23h). Among the tissues that give rise to injury MSC, we validated these results specifically on the long bone. We chose *Mfap4* as a marker gene, as it is specifically expressed in injury MSCs in the long bone (Figure 6e, Supplementary figure ). Immunohistochemistry results showed that *Mfap4* is highly expressed in the long bone after injury, and is localized near the injury sites (Figure 6f). In contrast, *Mfap4* expression was rarely detected in uninjured long bone (Figure 6f).

Building on previous research that identified osteogenic potential in cells from injured tissues like skeletal muscle^62^ and calvarial bone^63^, we applied the **Differentiation Model** to the extended atlas (Figure 6a). This analysis revealed a distinct trajectory for injury-derived MSC OPCSTs transitioning towards osteoblasts, deviating from the trajectory of typical MSC OPCSTs (Figure 6g). We employed **TrajDiff** to explore the heterogeneity in the differentiation processes between these two populations. The cell abundance analysis suggested that injury-derived MSC OPCSTs exhibited higher density in the early stages of differentiation, but this density was lower in the late stages (Supplementary Figure 24a). This pattern was particularly evident in tissues like skeletal muscle, where cells in the late stages were scarce (Supplementary Figure 24a). These findings suggest that injury-derived MSCs might have a slower differentiation speed^52^ than typical MSCs.

Next, we performed a differential expression analysis using **TrajDiff**, which revealed distinct gene expression patterns. Genes associated with wound healing (e.g., *Mia3*, *Hif1a*, *Smoc2*) and chemotaxis (e.g., *Cxcl9*, *Lrp1*, *Ecscr*) were consistently upregulated throughout differentiation in injury-derived MSCs (*Up*-*Up*) (Figure 6h,i, Supplementary Figure 24b). Conversely, genes involved in cell fate commitment (e.g., *Hes1*, *Bcl11b*, *Id2*,) were downregulated (Down-Down) (Figure 6h,i, Supplementary Figure 24b). This suggests that injury may steer these cells away from terminal differentiation states. Additionally, genes related to interleukin-1 beta production (e.g., *Cd36*, *Ccl3*, *Igf1*) showed early upregulation (*Up_0*), indicating a potential role in the initial response to injury (Figure 6h,i, Supplementary Figure 24b). Interestingly, genes involved in growth factor activity (e.g., *Ogn*, *Hbegf*, *Clec11a*) displayed a transient upregulation pattern, being upregulated early but downregulated later, suggesting a stage-specific role for growth factor activity in injury MSC differentiation (Figure 6h,i, Supplementary Figure 24b).

We then applied **TRAVMap** to our extended atlas (Supplementary Figure 25a-f). This analysis identified three injury-related gene modules (Supplementary Figure 25g-k). These modules exhibited high activity specifically in injury trajectories, primarily functioning at the early stage of differentiation (Supplementary Figure 25k). GO enrichment analysis revealed that these modules (TRAV540, TRAV766, TRAV767) are associated with distinct processes: immune response (*Sema7a*, *Rsad2*, *Tek*), bone development (*Scx*, *Acp5*, *Fam20c*), and cell proliferation (*Klf4*, *Egr1*, *Hpgd*), respectively (Supplementary Figure 25m).

## Discussion

In this study, we introduce **TrajAtlas**, a novel framework centered on trajectories to analyze differentiation heterogeneity. Unlike previous methods, **TrajAtlas** prioritizes trajectories to build a comprehensive landscape of osteoblast differentiation, representing the first framework to systematically dissect large-scale trajectory heterogeneity. **TrajAtlas** allows multi-scale exploration of differentiation, examining both individual cells and entire samples. This deepens our understanding of stem cell heterogeneity and how trajectories influence cell density, gene expression, and gene modules. Our key technical innovations include: (1) integrating large-scale trajectories to construct a reference atlas and a universal differentiation model, (2) developing statistical tools to identify differentially expressed genes along trajectories, and (3) implementing methods to infer conserved pseudotemporal gene modules across population-level samples. Additionally, we established methods to visualize changes in gene expression and gene modules across population-level datasets. To guarantee the framework’s universality for osteogenic processes, we initially built a core reference atlas using data from three tissues. We then successfully validated the framework on the extended atlas with various osteogenic datasets.

The major challenge in skeletal biology lies in the diversity of osteoprogenitor cells^5 6^. Due to a lack of comprehensive annotation, the characterization of responding trajectories has been a subject of controversy for a long time^5 6 7 30 39^. Our work addressed this gap by mapping experimentally validated osteoprogenitors onto our detailed differentiation atlas, which utilizes a seven-level annotation system. This analysis categorized osteoprogenitors into six major clusters, with at least four exhibiting the potential to directly transform into osteoblasts. We further identified key transcription factor groups that differentially regulate this transformation process, termed osteoprogenitor cell-specific differentiation trajectories. By investigating several crucial pathways for osteoblast differentiation, we revealed distinct activity patterns during osteogenesis, influenced by both the cellular source of the osteoprogenitors and the stage of differentiation. For example, the Hedgehog signaling pathway, known for its role in bone formation and maintenance^64^, displayed varying activity levels during the early stages of MSC OPCSTs, with *Hhip* and *Tubb6* identified as potential target genes. This OPCST-dependent pathway activity suggested an osteoprogneitor-specific approach for selecting therapeutic targets for bone diseases^65^. Finally, by applying **TRAVMap**, we identified dozens of pseudotemporal gene modules that function sequentially throughout OPCST differentiation. This work sheds light on the dynamic interplay between genes and gene modules as differentiation progresses.

Bone formation and bone disease are closely related with age^16, 39^ and tissue location^5 6^. In this study, we investigated the influence of age and tissue location on bone formation and disease, focusing on four key aspects: cell type composition, osteoprogenitor state, differentially expressed genes along osteoprogenitors’ differentiation trajectories and gene modules. We identified universal age-related hallmarks across osteoprogenitors, such as cell cycle activity and Wnt pathway activation, which aligned with previous findings on senescence^41^. **TRAVMap** analysis revealed a similar conserved effect in gene modules, consistently associating specific modules with age across multiple OPCSTs. This age-related influence was likely mediated by *Fosl1*(+), suggesting potential therapeutic targets for age-related bone diseases like osteoporosis. Notably, **TrajDiff** analysis of LepR+ BMSC OPCSTs revealed differentially expressed genes categorized by their expression patterns throughout differentiation. This finding highlights the highly specific and variable effect of age on different OPCSTs and differentiation stages.

Several tissues are widely thought to contribute to bone regeneration, including bone marrow^7^, periosteum^40^, and skeletal muscle^62^. By analyzing datasets from injured tissues, we identified an injury-related MSC-like cellular state. This phenomenon resembled blastema formation observed in slamanders^60^ and fish^66^, where blastemas arise from the dermis, cartilage, and muscle cells. Interestingly, mature osteoblasts were only observed in datasets derived from long bones and calvarial bones, suggesting a potential loss of regenerative capacity in mammals^67^. Compared to typical MSCs, injury MSCs exhibited altered differentiation trajectories. Using **TrajDiff**, we identified genes and gene modules associated with injury, providing valuable insights into bone regeneration processes.

While our framework is primarily designed to analyze osteoblast differentiation, it has the potential to be adapted for studying other differentiation processes, cell reprogramming, and even the cell cycle. Traditional “cell-centric analysis” offers a snapshot of cell type and gene expression at a single point in time. In contrast, “trajectory-centric analysis” sheds light on dynamic processes by examining changes over time. In summary, our study introduces a novel trajectory-centric framework that provides new insights into the dynamic interplay between cells, genes, and gene modules during osteogenesis.

## Materials and Methods

### Dataset collection

#### Data source

All datasets were derived from published studies. In **Differentiation Atlas**, to establish a representative landscape of osteoblast differentiation, datasets satisfied the following requirements were included: 1. Samples were collected from three tissue origins: the head, limb buds, and long bones, which serve as the primary sources of osteoblasts in mice^5 6^. 2. Samples were collected from healthy mice without additional treatment. 3. Representative of a specific aspect of osteoblast formation. For example, GSE190616^33^ was selected for inclusion due to its derivation from *Col10a1*-cre mice, which represents a pivotal transformation from hyperchondrocytes to osteoblasts. After selection, the **Differentiation Atlas** comprises datasets from 26 studies (Supplementary Figure 1a-c, Supplementary table 1).

Single-cell datasets related to osteogenesis were included in the extended atlas, to capture the diversity of osteogenesis. The extended atlas encompasses a broader range of tissues, including digit, rib bone, periodontium, and skeletal muscle; a wider array of treatments, such as disease models, drug interventions, and gene knockdown experiments; and multiple species, including human and rat models. Finally, 66 studies were included in the extended datasets (Supplementary Figure 22a-f, Supplementary table 1).

#### Metadata collection

For each study sample, we collected metadata, including information on the age, tissue origin, sequencing method, species, genotype, treatment methods, tissue dissociation methods, associated literature, and GEO accession (Supplementary table 1). We divided age into six groups, based on previous standards and osteogenesis changes. Organogenesis stage (E8.5-E14)^68^, Fetal stage (E14.5-E18.5), Postnatal (P0-P30), Young adult (1M-3M)^39^, Adult (3M-18M), Old (>12M)^69^. We divided tissue origin into head, limb bud ,and long bone.

According to previous studies, these three tissues represent three different sources of osteoblasts. Specifically, osteoblasts in the head region are derived from mesenchyme and fibroblasts, those in the limb bud from perichondrium and chondrocytes, and those in the long bone from chondrocytes and BMSC^5 6^. The detailed description was provided in supplementary note 1.

### Preparation of differentiation atlas construction

#### Data preprocessing

We performed two rounds of preprocessing.

In the first round of preprocessing, our objective was to filter out cells unrelated to osteogenesis and conduct annotation for integration in every dataset separately for each dataset. In detail, we excluded cells with fewer than 300 RNA features and fewer than 800 RNA counts. Additionally, cells with a high proportion (>20%) of transcript counts derived from mitochondrial-encoded genes were removed. Genes present in fewer than 3 cells in the datasets were also eliminated. We utilized **Seurat**’s normalization method, LogNormalize, with a scale factor set to 10,000. The FindVariableFeatures function in **Seurat** was employed to select the top 2000 highly variable genes for downstream analysis. Principal Component Analysis (PCA) was applied for dimensionality reduction, considering the top 30 dimensions for analysis. For studies involving multiple samples, batch effects were corrected using the **Harmony** algorithm^70^. We employed the **Louvain** algorithm for constructing a shared nearest-neighbor graph and conducting clustering analysis, implemented in the FindNeighbors and FindClusters functions. Subsequently, we visualized the identified clusters using the UMAP method. We performed two types of annotations. The first annotation was based on the study from which the datasets originated. The second annotation was conducted based on prior knowledge, manually assigning cells to eight categories based on marker genes (Supplementary Note 2). After the annotation process, we excluded cell types that were not related to osteogenesis, such as epithelial cells, from further analysis. Cell clusters meeting either of the following criteria were considered relevant to osteogenesis and included in the differentiation atlas: 1. Literature reports indicating a clear ability to form osteoblasts, such as chondrocytes. 2. Continuous adjacency with osteoblast cells on the reduction of diffusion map or UMAP.

The second round of preprocessing included droplet detection, count normalization, and highly variable gene selection. Specifically, we employed **scDblFinder**^71^ for droplet detection, removing cell barcodes marked as doublets from the matrix. To address differences in total UMI counts per cell, we conducted **SCRAN** normalization, following the procedure described in the Human Lung Cell Atlas (HLCA)^22^. The process involved total count normalization, log transformation, PCA, neighborhood graph calculation, **Louvain** clustering, and subsequent **SCRAN** normalization on raw counts using **Louvain** clusters. The resulting size factors from this process were utilized for normalization. For the final datasets, cells with unusually low size factors or exceptionally high total counts after normalization were excluded from the data. Finally, we made highly variable genes selection by **Scanpy**’s highly_variable_genes function^72^.

#### Batch division

Before integration, deciding which datasets should be divided into batches is crucial to preserve biological variance and eliminate technical noise. We employed **kBET**^73^, a tool based on k-nearest neighbors to quantify batch effects and determine which studies should be split into multiple batches.

#### Integration Method Benchmarking and Parameter Selection

The effectiveness of integration relies on several factors, including the integration tools, data normalization methods, the selection of highly variable genes, parameters of the integration tool, and cell annotation (especially for integration algorithms dependent on cell type information). To choose the optimal integration method for the Differentiation Atlas, we conducted benchmarking on the core datasets using the **sciB-pipeline** benchmarking framework^74^.

For integration methods, we considered 12 alternatives, including **Scanorama**^75^, **Harmony**^70^, **bbknn**^76^, **scANVI**^29^, among others. Annotation categories were classified as complete annotation, no annotation, and incomplete annotation (Osteoblast and Non-osteoblast).

Therefore, we performed a total of nineteen combinations, considering data normalization, integration tools, and annotation methods (Supplementary Figure 2a). Finally, we utilized **scib-metric** to measure the extent of batch effect removal and preservation of biological variation for each combination. In the end, we opted for **scANVI** as the final integration solution due to its high performance score (Supplementary Figure 2a).

We employed the scvi.autotune.ModelTuner function to select hyperparameters, including n_hidden and gene_likelihood. Additionally, **scib-metrics** was utilized to determine the optimal hyperparameters for the number of variable genes and n_latent (Supplementary Figure 2b).

### Construction of Differentiation Atlas

#### Integration of differentiation atlas with scANVI

For the integration of datasets into **Differentiation Atlas**, incompleted annotation (Osteoblast, Non-osteoblast) was used for integration benchmarking. **scANVI** was run on raw counts of 1500 highly variable genes (HVGs) (Supplementary Figure 2b). The batch variable was described in *batch division*. While running **scANVI**, the following parameters were used: n_latent:1500, encode_covariates: True, use_layer_norm: both, use_batch_norm: none, gene_likelihood: zinb, n_layers:1, dropout_rate: 0.1,max_epochs: 100 (Supplementary Figure 2b).

#### Cluster detection and annotation

In general, we employed **scHarmonization** pipeline^23^ (https://github.com/lsteuernagel/scHarmonization) for multi-level clustering. Initially, we applied the **Leiden** algorithm for clustering, incrementing the clustering resolution from 0.001 to 50. The Leiden clustering results with progressively increasing resolutions were selected, aiming to approximately double the number of clusters at each level. The **ROGUE** package^77^ was used to evaluate the clustering results and the selection of the clustering cutoff values.

The first level of clustering typically distinguishes between osteoblast and non-osteoblast cells, while the second level aims to differentiate major cell types such as chondrocytes and MSCs, consistent with coarse label hierarchy. For the subsequent five levels, the **mrtree** algorithm^23^ was used to form a clustering tree. The marker genes were selected using specificity scores as described in the **scHarmonization** pipeline^23^. After determining the marker genes (specificity > 1) for each cluster node in the tree, sibling clusters with fewer than 10 related markers were merged into a single cluster node to avoid over-clustering. Visualization of the clustering tree was performed using the **ggtree** R package. The van Elteren test implementation for **Seurat** objects (https://github.com/KChen-lab/stratified-tests-for-seurat) was utilized for detecting marker genes.

We manually assigned the first three levels of annotation based on tissue origins and cell markers of cell clusters. For clusters in higher levels, we concatenated the node’s best marker gene with its parent’s name. The details of the seven-level annotation can be interactively explored at https://zyflab.shinyapps.io/TrajAtlas_shiny/.

#### Osteoprogenitor Cells Mapping

We collected data on experimentally validated osteoprogenitor cells (EV-OPCs) using either transplantation assays or Cre-loxP system-driven lineage tracing strategies^6^ from a total of 28 studies. The collected details encompassed cell markers, publication years, study names, journals, and tissue location (Supplementary table 3). The cell clusters in our differentiation atlas exhibiting high expression of EV-OPC markers, and corresponding to tissues where EV-OPCs were located, were considered to be accurately mapped. Most EV-OPCs (24/28) can be mapped to our differentiation atlas. The details of OPCs Mapping can be interactively explored at https://zyflab.shinyapps.io/TrajAtlas_shiny/.

#### Covariate Impact on Osteoprogenitor Cells

We split covariates as technical factors (UMI per cell, number of genes detected) and biological factors (Age, Tissue) (Supplementary Figure 6a). We utilized the correlation between covariates and variation in the atlas to depict how covariates influence cellular states^22^. We employed principal component regression on each covariate with scANVI latent component scores, as described in ref.^22^.

Following, we used the **dreamlet** package^78^ to perform differential expression analysis, which applied a linear mixed model to sample-level pseudobulk. To identify age-correlated differential genes, we encoded age covariates numerically, with ’Organogenesis stage’ assigned a value of 1 and ’Old’ assigned a value of 6. We set “∼Age” as the design formula to identify genes linearly correlated with age covariates. Genes that exhibited downregulation across all osteoprogenitors were categorized as “concordant downregulated genes”, while those showing upregulation across all osteoprogenitors were termed “concordant upregulated genes”. Genes that did not follow either pattern, showing mixed regulation across osteoprogenitors , were labeled as “divergently regulated genes” (Supplementary Figure 6b,c). Gene set analysis was performed with **zenith** package (https://bioconductor.org/packages/release/bioc/html/zenith.html) with M2 gene sets in **MisgDB**.

### Lineage and pseudotime inference

#### Differentiation model construction

The construction of the **Differentiation Model** encompassed the detection of transition probabilities, identification of endpoints, inference of paths, reconstruction of pseudotime, construction of lineage models, inference of GRNs, and analysis of genes and pathways.

We utilized **PAGA**^28^ to measure coarse-grained transition probabilities between osteoprogenitors and osteoblasts. PAGA scores greater than 0.015 were defined as connected, while scores below this threshold were considered not connected.

For endpoint identification, we utilized prior knowledge, **CytoTRACE**, development time point of the samples to identify the endpoint of OPCST trajectory (Supplementary Note 3).

Subsequently, we inferred the lineage path by **Slingshot**^79^. The pseudotime for each OPCST was individually inferred using **Palantir**^80^. To ensure the comparability of pseudotime across different OPCST, we employed the generalization capability of machine learning techniques (Supplementary Note 4). We utilized the **LightGBM** model due to its demonstrated excellent accuracy and generalization ability compared to other models (Supplementary Figure 9b,c, Supplementary note 4). We have shown that this model can be transferred to other single-cell datasets or bulk datasets to accurately predict the progression of osteoblast differentiation (Supplementary Figure 9f-h, Supplementary Note 4).

To construct the lineage model, we partitioned each OPCST into 10 bins based on pseudotime, ensuring that each bin had an equal pseudotime gap. Subsequently, if bins at the same pseudotime across different OPCSTs exhibited a high similarity score, we merged those bins. The similarity measurement was conducted using a random forest model by assessing whether one bin could predict the other. In detail, we generated reference latent representations from the same cells sampled from each bin to train a random forest classifier. Utilizing misclassification rates, we computed a similarity score, reflecting the degree of resemblance between the bins.

We utilized **SCENIC**^24^ to infer the transcription factor activity of each bin. We annotated each transcription factor, indicating whether it has been reported to be associated with osteogenesis (Supplementary table 5). We utilized **tradeSeq** to build a GAM model to model pseudotemporal gene expression, then clustered genes into 12 clusters with K-means. Gene set enrichment was performed with **clusterProfiler**^81^ with Reactome database. The pathway activity was inferred by **AUCell** (https://github.com/aertslab/AUCell).

To construct a comprehensive osteogenesis gene database, we combined differential genes in our model with the Phylobone database^42^, bone-related GWAS SNPs^44^, bone-related genes from Gene Ontology (GO) terms^44^, and bone signature genes from other publications^43 44^ to annotate bone-related genes.

For the trajectory dotplot, we extracted three attributes from trajectory: correlation, peak, and expression (Supplementary Note 5). For “correlation”, we calculated the Pearson correlation coefficient between gene expression and pseudotime. For “peak”, we divided pseudotime into 10 bins and used bins that had the max expression to represent the peak. For “expression”, we calculated the mean value of 10 bins as expr. We have proved that these three attributes can recover most information on expression patterns along pseudotime (Supplementary Note 5).

### Detecting differential abundance and expression along pseudotime (TrajDiff)

Both the differential abundance (DA) and differential expression (DE) follow this pipeline: neighborhood construction, neighborhoods differential test, and pseudotime-association test. The benchmarking of **TrajDiff** and other differential pseudotime analysis tools (**Condiments**, **Lamian**) was described in Supplementary Note 6.

#### Differential abundance

The first two steps, neighborhood construction, and local differential testing followed the methods described in **MiloR**^82^. In short, we constructed a KNN graph and defined cell neighborhoods in the neighborhood construction step. In the local differential testing step, we counted cells in the neighborhoods to construct an *N* × *S* (neighborhood × experimental sample count) matrix, then used **edgeR** to perform differential analysis, followed by controlling the spatial FDR in neighborhoods utilizing methods described in **Cydar**^83^. In the pseudotime-association test, we projected each neighborhood on a pseudotime axis. For neighborhoods with spatial FDR below a threshold (0.05 by default), we labeled them as “Rejection”; otherwise, we labeled them as “Accept”. Then we divided the pseudotime axis into *n* intervals (100 by default) and calculated the counts of Accept (N_accept_) and Rejection (N_rejection_) for each interval. Then we used binomial distribution to test whether each interval had a difference as follows:

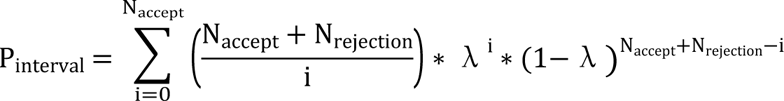

For λ in this equation, we generated null datasets by randomly shuffling the order of samples. λ was then calculated as 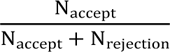 for the null datasets. We repeated this process multiple times (*n*) to calculate the average value of λ and improve precision. If 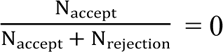, we set λ to the precision of the calculation, i.e., 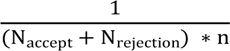.

For overall *p-*value(P_overall_), we replaced N_accept_ and N_rejection_ to the count of Accept and Rejection of the whole neighborhood.

We computed the mean count per million (*CPM*) within each interval derived by **edgeR** to generate CPM_interval_, representing the model-fitted cell abundance. We quantified the difference in cell abundance by multiplying the log-fold change (*logFC*) and *CPM* calculated by **edgeR**, and calculated the mean value for each interval as follows:

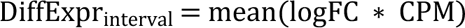

#### Differential expression

Firstly we constructed neighborhoods as described in the section *Differential abundance*. In the neighborhoods differential test step, we made a pseudobulk for each neighborhood and constructed an *N* × *S* × *G* (neighborhood × experimental sample count × gene count) matrix. Then we used **edgeR** to loop every gene to perform differential analysis, followed by controlling the spatial FDR.

In the pseudotime-association test, we employed the identical procedure outlined in the description of the *differential abundance*. We iterated through each gene to generate P_interval_, CPM_interval_ and DiffExpr_interval_, representing the significance of differential expression, model-fitted expression, and model-fitted expression difference, respectively. GO enrichment analysis was conducted using **clusterProfiler**. For GO-enriched genes, we used trajectory dotplots to visualize their pseudotemporal expression patterns across trajectories.

### Construction of TRAVMap

The TRAVs were identified through matrix factorization of the pseudotemporal expression matrix derived from trajectories. In detail, For each trajectory defined in **Differentiation Model** construction, we derived CPM_interval_ with **TrajDiff** to form a *G* × 100 (gene count × interval count) matrix. We filtered out empty intervals and selected the top 2000 highly variable genes (refer to *Data preprocessing*), followed by scaling matrix by row. Then we applied non-negative matrix factorization (NMF)^54^ for matrix factorization, decomposing each dataset into 15 factors (the number of factors was chosen by cross-validation). Merging all factors, we performed hierarchical clustering^55^ to form 226 clusters (=round (datasets count × factor count) / 8). We calculated the mean signal of factors in each cluster denoted as TRAVs. We extracted the top 100 genes to define the TRAV gene modules. We referred to the methodology described in ref.^55^ and calculated the correlation coefficient between TRAVs and the NMF factors of each trajectory, denoting it as TRAV activity. The gene expression patterns of gene modules were visualized with trajectory dotplots. We calculated the expression, peak, and correlation in the trajectories with the highest TRAV activity to represent the expression pattern of each gene module. The GO enrichment of TRAV gene modules was conducted by **clusterProfiler**. The transcription factors of TRAV gene modules were predicted using the TRRUST database via **Enrichr** (https://maayanlab.cloud/Enrichr/).

To visualize gene module activity on population-level trajectories, we utilized TRAV activities and gene expression patterns to learn trajectory representations. The detailed methods for trajectory reduction were described in the Supplementary Note 7.

We used the FindMarkers function in **Seurat** to detect OPCST-related TRAVs, denoting TRAVs that have high activity (0.4) in more than one-third of trajectories as osteogenesis-conserved TRAVs. We utilized a linear mixed model to detect age-related TRAVs (C*ovariate Impact on Osteoprogenitors* ).

#### Mice

Female C57BL/6 mice were obtained from the Animal Center of Wuhan University. All experimental procedures were approved by Wuhan University and were performed according to laboratory animal care and use guidelines. The study protocol was approved by the Ethics Committee for Animal Use of the Institute of Biomedical Sciences (Protocol number 69/2017).

For the injury model, female C57BL/6 mice (8 weeks old) were operated on for ablation surgery. Right femurs were operated, while left femurs were untreated and used as an internal control. We followed the procedure described in the previous study^7^. To create a bone marrow injury model, an incision was made on the skin, the knee ligaments were separated, and a cylindrical area of the marrow space in the femur was sequentially removed using endodontic instruments of increasing gauge sizes. The surgical site was irrigated with saline, and the incision was sutured closed.

### Histology and immunohistochemistry

For histochemical analysis, the femurs were fixed in 4% paraformaldehyde at 4°C for 24 hours, decalcified in 10% EDTA (pH 7.4) for 4 weeks, and subsequently embedded in paraffin. 5 μm sections were prepared using a microtome (Leica). For immunohistochemical staining, the sections were digested with 0.05% trypsin at 37°C for 15 minutes. The sections were then incubated with anti-MFAP4 antibody (1:200; ThermoFisher) overnight at 4°C, followed by incubation with goat anti-rabbit Alexa Fluor 488 secondary antibody. Fluorescence was visualized using a fluorescence microscope (Thunder Imager; Leica Microsystems).

## Data availability

The **Differentiation Atlas** (raw counts, integrated embedding, cell type annotations, clinical and technical metadata) is publicly available and can be downloaded via Figshare (https://figshare.com/articles/dataset/Differential_Atlas/25422688). The annotation system and osteoprogenitor mapping of **Differentiation Atlas** can be interactively explored at https://zyflab.shinyapps.io/TrajAtlas_shiny/.

## Codes availability

The **TrajAtlas** software package is available at https://github.com/GilbertHan1011/TrajAtlas. The **TrajAtlas** documentation including API, tutorials ,and examples is available at https://trajatlas.readthedocs.io/. Codes to reproduce our analysis and figures are available at https://github.com/GilbertHan1011/TrajAtlasManuscript.

